# Unique pattern of injuries in *Olenoides serratus* from the Burgess Shale elucidate moulting process in trilobites

**DOI:** 10.64898/2026.09.18.752702

**Authors:** Sarah R. Losso, Russell D. C. Bicknell

## Abstract

Ecdysozoans is one of the most abundant and diverse groups of metazoans and grow through moulting the rigid exoskeleton that provided protection. But the moulting process is often dangerous, resulting in malformations or death. Injuries have been well documented in the calcitic exoskeleton of trilobites, some of which have been attributed to moulting complications when long spines are deformed. Injuries in trilobites are often found in the trunk rather than the cephalon, and abnormal genal spines are rarely documented. Here we examine injuries in eleven specimens of *Olenoides serratus* from the Burgess Shale (Cambrian, Wuluian) and demonstrate an increased frequency of genal spine malformations compared to other trilobites. The unusual pattern of malformations may be attributed to moulting injuries of the long and delicate genal spines in *O. serratus*. To moult, trilobites curved ventrally to press the anterior margin of their cephalon into the substrate and open the facial sutures. This process exerted force on the librigenae. To exit the exuvia, the genal spines were forced to bend as the old librigenae dipped ventrally along the anterior edge, resulting in the injured genal spines in *O. serratus*. A similar process of moulting resulting in injuries may be seen in other early Euarthropods.

## Introduction

Moulting is a crucial process within ecdysozoans, allowing animals to grow in stages while maintaining the protection provided by the tough exoskeleton. Ecdysozoans are some of the most abundant and diverse groups of animals and have been since the Cambrian (Budd and Jensen, 2000; Daley and Drage, 2016). Despite the benefits of moulting, it is a dangerous process (Bowser and Rosemark, 1981; Fujaya et al., 2013). Successful moulting (i.e. survival) in marine euarthropods depends on a variety of factors including temperature, salinity, food availability and nutrition (Bowser and Rosemark, 1981; Gong et al., 2015; Noordin et al., 2020). Despite a successful moult, the individual may be malformed. Moulting injuries and the vulnerability during the soft post-moult phase have been suggested to result in up to 90% of mortality in modern euarthropods (Clarkson, 1979).

Trilobites are an abundant and diverse group of Paleozoic euarthropods due to their readily fossilized calcite dorsal exoskeleton. Malformations in the exoskeleton have been documented in many species of trilobites and have been attributed to injury, genetic or embryological malfunctions, or pathology (Owen, 1985; Babcock, 2007). While malformations in trilobites are well documented, they are commonly only known from a singular specimen of a species (Bicknell et al., 2024, 2025, 2026). Injuries can occur through predation, intraspecific competition and through accidental damage including moulting (Owen, 1985; Babcock, 1993) and are most commonly reported from larger specimens (Bicknell et al., 2022a, 2024). Examination injury stereotypy has demonstrated a bias towards posteriorly located malformations (Babcock, 1993; Pates et al., 2017; Zong et al., 2023; Bicknell et al., 2024; Mahata and Pates, 2026).

*Olenoides serratus* a well-documented trilobite species, with abundant specimens and appendage preservation (Whittington, 1975, 1980; Losso et al., 2025). Several instances of injuries have been reported due to the abundance of the species (Babcock, 1993, 2007; Pratt, 1998; Bicknell and Paterson, 2018; Bicknell et al., 2024). However, patterns of size, injury stereotypy and possible cause have not extensively been explored due to a limited number of malformed specimens and lack of examination within this framework.

## Materials and Methods

### Specimens

All studied specimens are housed at the invertebrate paleontology collections at the Smithsonian Institution (USNM) Washington, D.C., USA, the Geological Survey of Canada (GSC) Ottawa, Ontario, Canada, at the Royal Ontario Museum (ROM) Toronto, Ontario, Canada, and Naturmuseum Senckenberg (SMF) Frankfurt, Germany. For additional institutional abbreviations see Supplemental Table 1.

### Photography

Specimens were photographed under cross polarized light using a Nikon D850 DSLR camera fitted with a Macro Nikkor 60 mm lens, a Nikon D7500 DSLR camera fitted with a Macro Nikkor 40 mm lens, or Olympus E-M1 MarkIII fitted with a 60 mm macro lens.

### Measurements

Glabellar length and width at the L3 were measured in all specimens. Malformations were identified with location, left or right side, and type of injury (Supplemental Table 2). Graphs of frequency injuries and distribution through size were produced in R (R Core Team, 2017).

### Comparative assessments

Eight trilobite species (*Olenoides serratus*, *Eccaparadoxides pradoanus, Paradoxides davidis trapezopyge, Redlichia takooensis, Redlichia rex, Ogygopsis klotzi, Ogygiocarella debuchii,* and *Arctinurus boltoni*) have been documented with more than three injured specimens. For these species, each injury location was collated and divided into 12 zones across the exoskeleton (Supplemental Fig. 1; Table 3). Where multiple injuries are noted on a same specimen, all injuries were recorded independently (Supplemental Table 3). Histograms of the distribution injuries across all 12 zones were produced for each species (Supplemental Fig. 2) and used to illustrate most frequently damaged areas on ling drawings of the exoskeletons.

## Results

### Descriptions of injuries

Five specimens have injuries to the left genal spine (Fig. 1).

**Figure 1.**
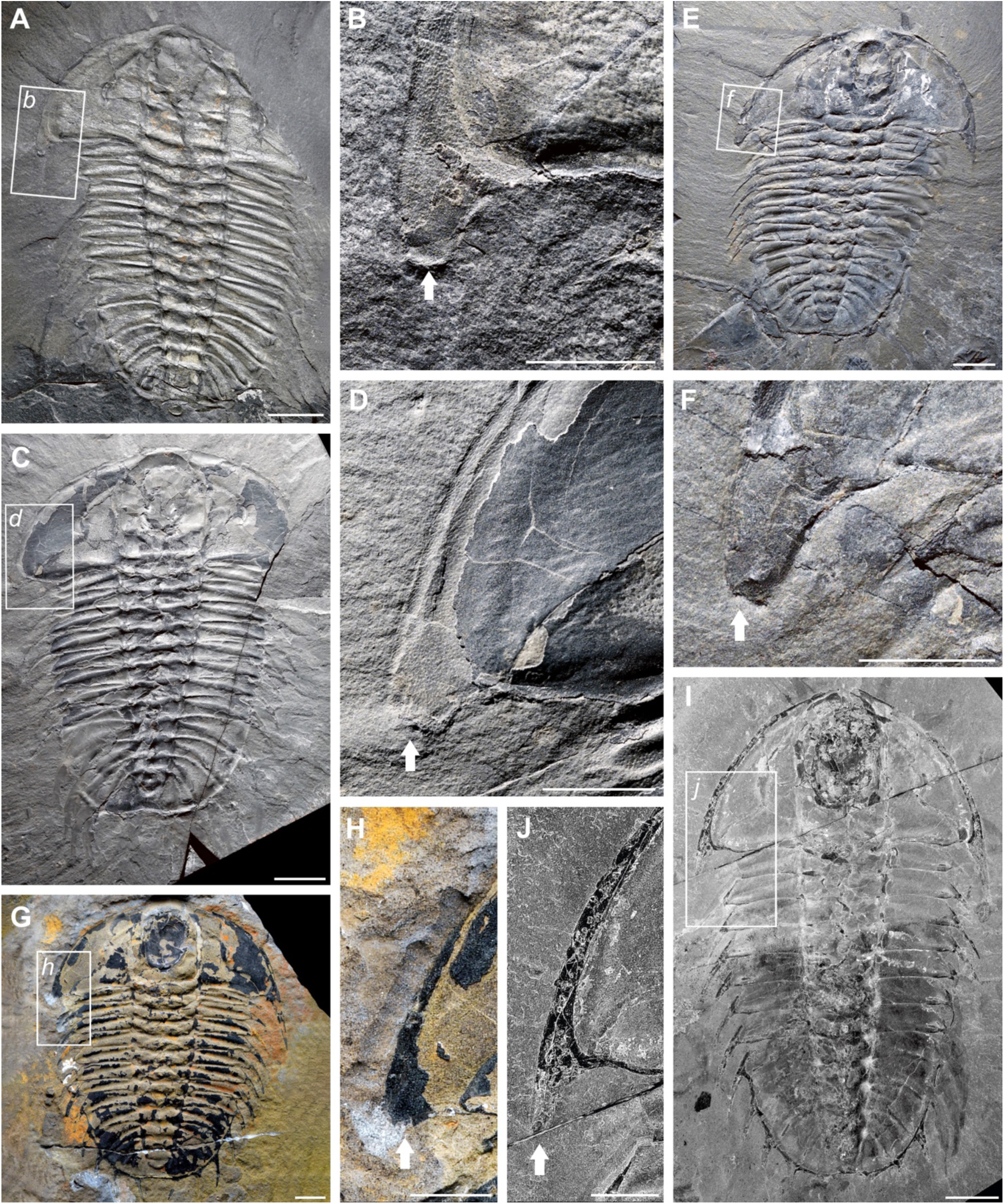
*Olenoides serratus* with genal spine injuries. (A, B): USMN 273253. (C, D) USMN 273257. (E, F) USNM 273324. (G, H): ROMIP 69734. (I, J) ROMIP 69733. Scale bars: (A, C, E, G, I): 10 mm; (B, D, F, H, J): 5 mm.

USMN 27325, a nearly complete specimen in dorsal view with the right librigenal lateral margin obscured by matrix, exhibits a truncated left genal spine (Fig. 1A). The left genal spine extends to the first thoracic tergite posterior margin. The posterior margin of the spine is rounded and slightly calloused (Fig. 1B).

USNM 273324 is a nearly complete specimen in ventral view with preserved pygidial appendages and has a damaged left genal spine (Fig. 1C). The right genal spine is injured and truncated, extending only to the first thoracic tergite intrapleural furrow (Fig. 1D). Posterior margin of the spine is concave (Fig. 1D).

USMN 273257, a complete specimen in dorsal view, exhibits a shortened left genal spine, extending to the second thoracic tergite intra pleural furrow (Fig. 1E, F).

ROMIP 69734 is a completely preserved specimen in dorsal view with a greatly reduced left genal spine (Fig. 1G). The left genal spine does not extend to the first thoracic tergite intrapleural furrow (Fig. 1H).

ROMIP 69733 is a complete specimen in dorsal view with preserved appendages (Fig. 1I). The left genal spine is shorter and more gracile than the right genal spine (Fig. 1I, J). The left genal spine extends to the interpleural furrow between thoracic tergites one and two (Fig. 1J).

Three specimens of *Olenoides serratus* exhibit injuries in the trunk (Fig. 2). ROMIP 14468 is a complete specimen preserved in dorsal view and exhibits two abnormalities on the left side (Fig. 2A). The left pleural spine of the seventh thoracic tergite is shortened with a blunt lateral margin (Fig. 2B). The pygidium shows a truncated and rounded posterior margin for the left third pygidial spine (Fig. 2A, C).

**Figure 2.**
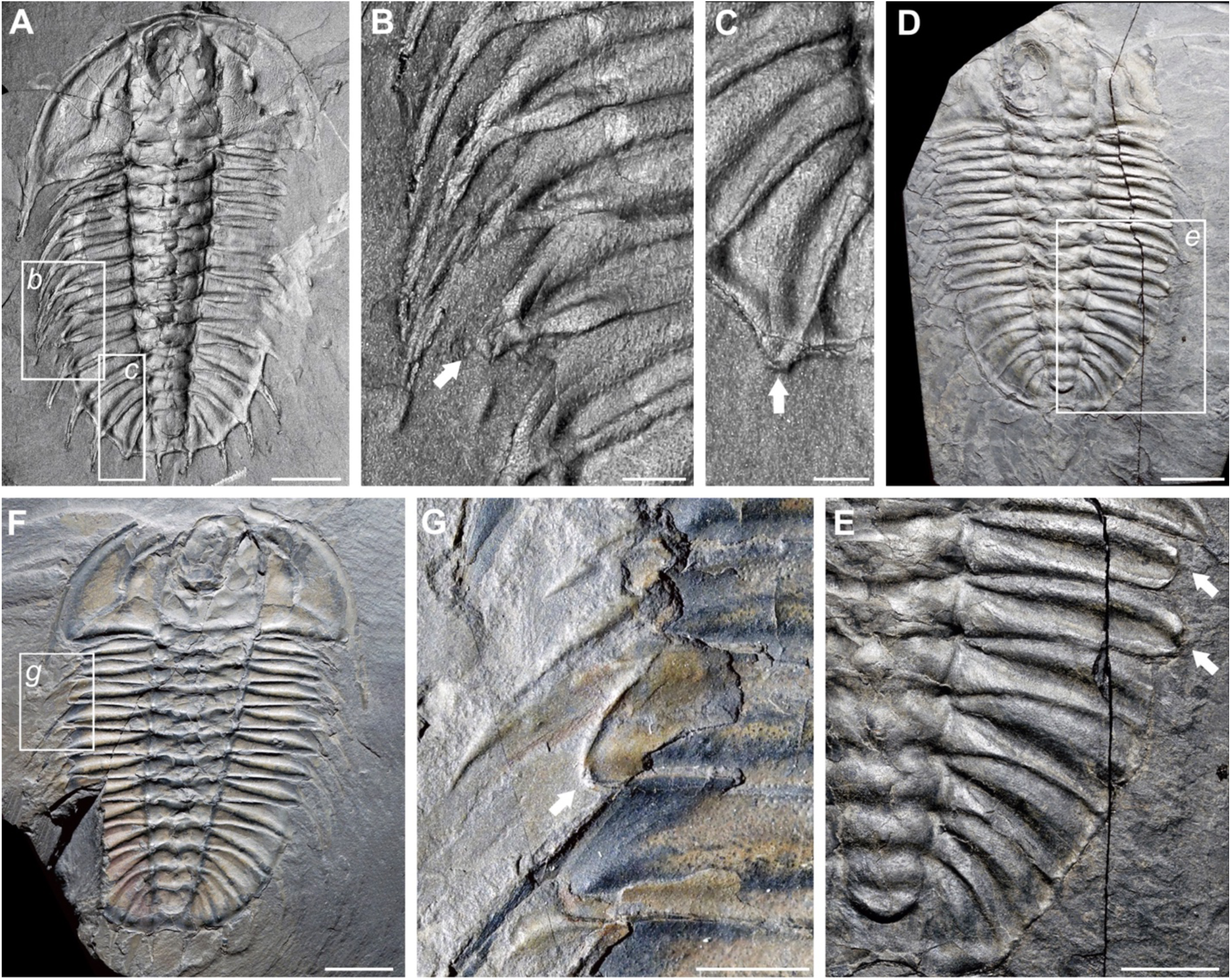
*Olenoides serratus* with thoracic and pygidial injuries. (A, B, C): ROMIP 14468. (D, E) USMN 274169. (F, G) USNM 273321.Scale bars: (A, D, F): 10 mm; (B, C, G): 2 mm; (E) 5 mm.

USMN 274169 is a nearly complete specimen, missing the left lateral and anterior margin on the cephalon, preserved in dorsal view (Fig. 2D). On the right side of USMN 274169, there is a ‘V’-shaped indentation at thoracic tergites six and seven that show blunt terminations and a complete lack of pleural spines (Fig. 2D, E).

USNM 273321 is a nearly complete ventrally preserved specimen (Fig. 2F). The right side of the third thoracic tergite exhibits a truncated and rounded pleural tip (Fig. 2G).

### Distribution of injuries

No specimens of *Olenoides serratus* have bilateral injuries. Of the 11 specimens identified with injuries, nine of them occur on the left side (Fig. 3A; Supplemental Table 1). A total of six specimens exhibit genal spine injuries, the five specimens in Fig. 1 and ROMIP 52497 figured by Pratt (1998) (Fig. 3B). Five genal spine injuries occur on the left side, and only one on the right (Figs. 1). Injuries in the trunk occur on the left side in four specimens, and right in one specimen (Figs. 2; 3; Supplemental Table 1).

**Figure 3.**
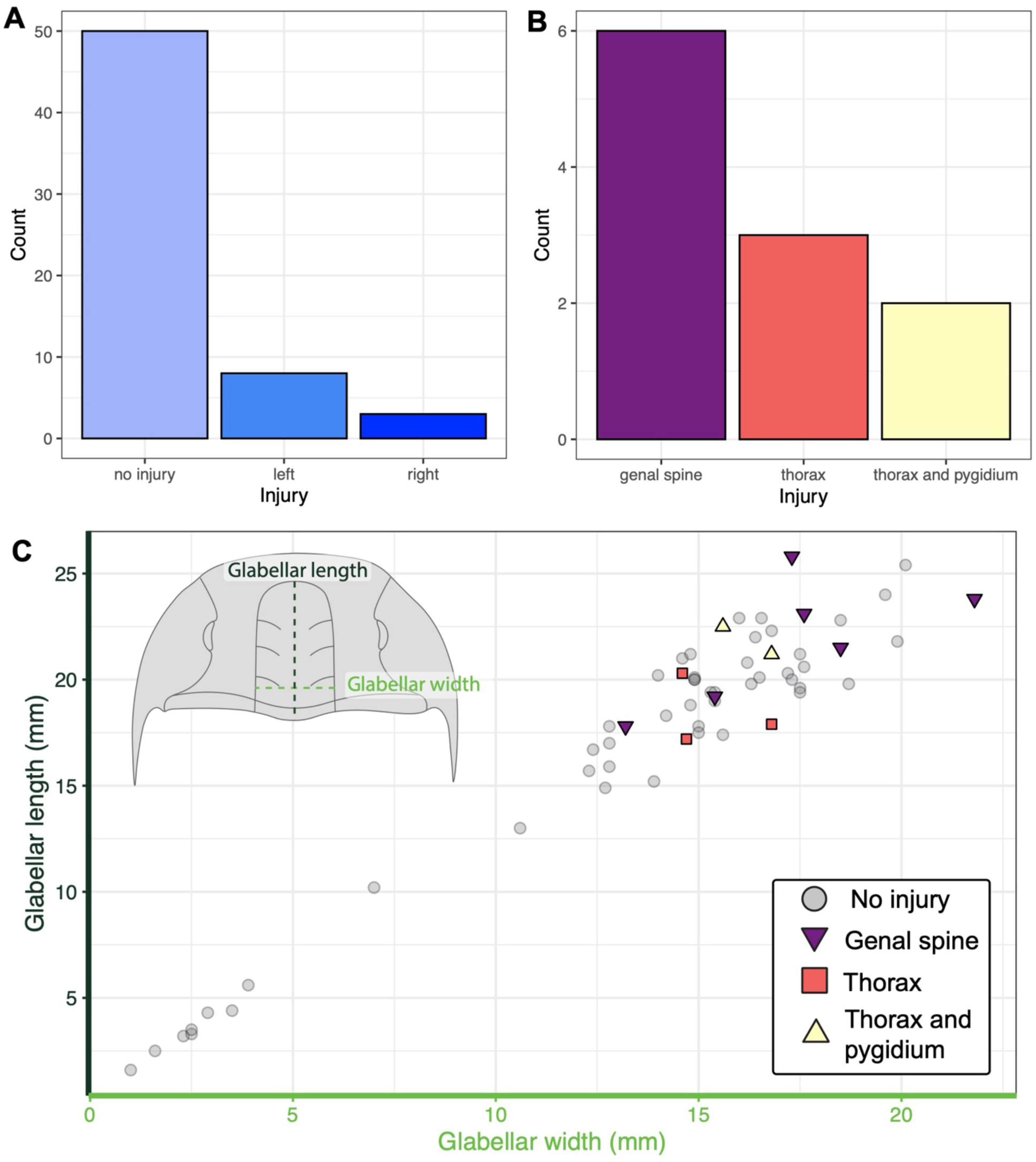
Distribution of injuries in *Olenoides serratus*. (A) Specimen size with injuries. (B) Distribution of injuries across tagma. (C) Distribution of injuries from left and right side of body.

Injuries have only been found in larger specimens (i.e. specimens larger than the median in glabellar length) (Fig. 3C; Supplemental Table 2, Supplemental Fig. 3). Genal spine injuries are found in a larger range of specimen sizes, from 13.2 – 21.8 mm in glabellar length (Fig. 3C). Injuries to the thorax or thorax in pygidium are found in specimens near the median size of glabellar length (Fig. 3C; Supplemental Table 2, Supplemental Fig. 3). Trunk injuries are found in specimens with glabella between 14.6 – 16.8 mm wide (Fig. 3C). Genal spine injuries are found in similarly sized specimens as those with trunk injuries, but also in large specimens (Fig. 3C). Two specimens with genal spine injuries — ROMIP 69734 and ROMIP 69733 —are some of the largest specimens of *Olenoides serratus* within the compiled dataset (Supplemental Table 1).

Trilobites with more than three injured individuals derive from the Cambrian (six species), and the Ordovician (one species) and the Silurian (one species) (Fig. 4). The posterior region of the thorax and pygidium is most frequently injured area in 75% of species (Fig. 4). In *Ogygopsis klotzi*, margins of thoracic tergites on the right side are most frequently injured. However, there is also evidence for injuries on the left thorax and anterior pygidium (Fig. 4F). Genal spine injuries are only seen in *Olenoides serratus* (Fig. 4a), *Ogygiocarella debuchii* (Fig. 4G), and *Arctinurus boltoni* (Fig. 4H).

**Figure 4.**
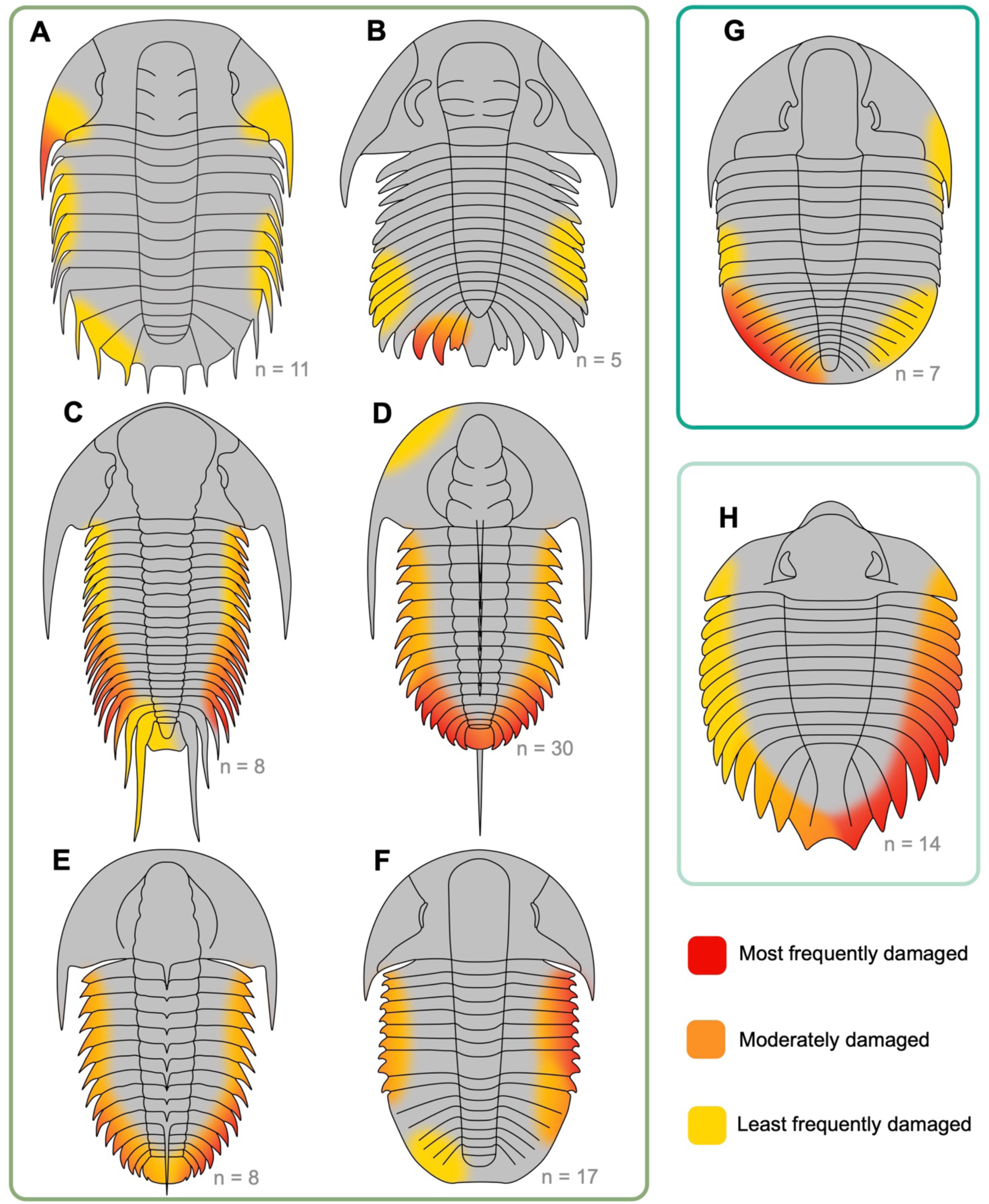
Occurrence of injuries in trilobites. (A) Olenoides serratus, this study. (B) Eccaparadoxides pradoanus. (C) Paradoxides davidis trapezopyge. (D) Redlichia takooensis. (E) Redlichia rex. (F) Ogygopsis klotzi. (G) Ogygiocarella debuchii. (H) Arctinurus boltoni. Forest green box denotes Cambrian taxa, dark green box denotes Ordovician taxon, and light green box denotes Silurian taxon. Numbers to the bottom right denote number of damaged specimens counted. In *Eccaparadoxides pradoanus* three additional specimens are figured in Pates et al. (2017) with hypostome or cephalic damage which were not considered here as those are not attributed to injury in life. See Supplemental Table 1 and Supplemental Figs. 1,2 for data.

## Discussion

### Size of injured specimens

Injuries are more frequently found in larger specimens within a single trilobite species. In *Redlichia rex* and *R. takooensis* from the Emu Bay Shale, injuries are only found in specimens that are above the median length (Bicknell et al., 2022a). *Elrathia kingii* from the Cambrian of the western USA shows a similar pattern of injuries restricted to the largest specimens (Bicknell et al., 2024). *Olenoides serratus* exhibits the same trend of injuries in large specimens (Fig. 3C). There may be a survivorship bias as larger specimens are more likely to survive attempted predation attacks, possibly from an increase in calcification, increased chance of non-lethal damage, or more opportunities for attacks from a longer lifespan (Bicknell et al., 2022a). In modern euarthropods, the cuticle protects the individual and requires a healthy microbiome, but the ability to repair injuries can decrease with age (Whitten et al., 2014; O’Neill et al., 2018; Liu et al., 2024). A similar process may have prevented older trilobites from repairing damage efficiently or make them more likely to incur damage.

### Distribution of injuries in trilobites

When taken together, injuries in trilobites most often occur on the right posterior side of the trunk (Babcock, 1993). However, more nuanced patterns are found when examining individual species or localities (Pates et al., 2017; Bicknell and Pates, 2020; Bicknell et al., 2022a; Zong et al., 2023; Mahata and Pates, 2026). In four species, damage is only known in trunk and is most common posteriorly without strong lateral bias (Fig. 4) as was reported in in trilobites from the Manuels River in Canada (Mahata and Pates, 2026). Cephalic injures are rarer than those in the trunk. Specimens of *Redlichia takooensis, Ogygiocarella debuchii* and *Arctinurus boltoni* are known with injured cephala, although in the former the anterior border is damaged rather than the genal spines (Fig. 4).

Damaged trilobite spines result in shorted morphologies with a calloused margin (Pates et al., 2017; Bicknell et al., 2024; Mahata and Pates, 2026). Pleural and pygidium spines are thought to be injured during moulting complications (Morris and Jenkins, 1985; Bicknell and Pates, 2020; Bicknell et al., 2022a, 2024). However, genal spines are seldom reported as injured. To date, only *Olenoides serratus* is known with multiple specimens displaying injuries to the genal spine.

Injuries in trilobites are generally rare despite the abundance of specimens throughout their 270-million-year history (Babcock, 1993). In the Cambrian, four species are known with more than three injured specimens, but this reduced throughout the Paleozoic and only one species is known from the Silurian Period. While there are additional instances of damage in fewer specimens (Bicknell and Smith, 2022; Zong et al., 2023). Injuries are apparently more commonly found in Cambrian and Ordovician trilobites, possibly resulting from changes in prey selection, genetics or sampling bias (Bicknell et al., 2024). However, this very much remains an open question.

### Potential causes of genal spine injuries

Injuries to the cephalon and axial lobe may be found rarely, likely because such damage would likely have been lethal (Babcock, 1993; Pratt, 1998; McNamara and Tuura, 2011; Mahata and Pates, 2026). The majority of vital organs in trilobites are concentrated with the axial lobe with only a thin membrane extending below the pleural lobes and librigenae (Lerosey-Aubril et al., 2012; Fatka et al., 2012; Gutiérrez-Marco et al., 2017; Kraft et al., 2023; El Albani et al., 2024; Losso and Ortega-Hernández, 2024). The genal spine of trilobites was used in some species to produce negative lift to allow the individual to remain on the seafloor (Pates and Drage, 2024), but it may not have been crucial to survival depending on the species lifestyle. In the modern Atlantic horseshoe crab, *Limulus polyphemus*, only injuries to structures with crucial functional roles (e.g. the telson used for right an overturned individual) are completely repaired (Bicknell and Pates, 2019; Bicknell et al., 2022b; Bicknell and Cuomo, 2024). As the genal spine injuries in *Olenoides serratus* were unlikely to be highly detrimental to the survival of an individual, the damage may have persisted through multiple moult cycles increasing the likelihood of preservation in the fossil record.

Injuries in trilobites may be caused by (1) predation, (2) teratologies, (3) pathologies or (4) accidental injury. Each cause may create distinct injuries or patterns of damage to the body and are here evaluated in turn as to potential causes of the damage in *Olenoides serratus*.

#### (1) Predation

Exoskeletal breakage that occurred when the animal was still alive because of failed predation. Injuries attributed to predation commonly have ‘L’, ‘U’, ‘V’, or ‘W’-shapes (Babcock, 1993; Bicknell et al., 2022a, 2023). These same morphologies often show evidence for cicatrization and repair, but this depends on the degree of exoskeletal recovery. Here, it seems likely that only USMN 274169 shows evidence of failed predation, given the truncation of multiple spines into a ‘V’-shaped indentation.

#### (2) Teratologies

These are external expressions of developmental, embryological, or genetic malfunctions observed on the exoskeleton (Owen, 1985; Bicknell et al., 2025). While rare, they can be associated with injuries. These morphologies include addition or removal of nodes, segments, and spines, as well as aberrant rib and furrow morphologies (Owen, 1985; Bicknell and Smith, 2022). We do not report any such malformations here.

#### (3) Pathologies

Malformed exoskeletal sections resulting from parasitic activity or infections. These structures are often expressed as circular to ovate swellings (Šnajdr, 1978; Owen, 1985; De Baets et al., 2021). We do not report any such malformations here.

#### (4) Accidental injury

Injuries derived from bumping a spine or difficulties moulting. Injuries from moulting reflect poor nutrients, stressful environments (Bowser and Rosemark, 1981; Gong et al., 2015; Noordin et al., 2020), or challenges exiting the exuvia. Within trilobites, injuries from moulting complications are commonly considered to be truncated, singular spines. Here, the majority of specimens we considered confirm to this condition. As such, USMN 27325, USNM 273324, USMN 273257, ROMIP 69734, ROMIP 69733, ROMIP 14468, and USNM 273321 all show evidence of accidental injury. The genal spine injuries are especially interesting as they present insight into how *Olenoides serratus* moulted.

Trilobites are thought to have arched into the sediment to apply force to the anterior margin of the cephalon and open the facial sutures to moult (Daley and Drage, 2016). *Olenoides serratus* had a 4-step moulting process: 1) facial sutures open; 2) rostral plate removed; 3) cephalon removed; 4) cranidium removed (Chen et al., 2008, 2023; Daley and Drage, 2016). As the facial sutures open and the librigenae are separated from the cranidium, the genal spines would be one of the first structures of the new moult freed from the exuvia. Although short trilobite spines likely undamaged during moulting (Conway Morris and Jenkins, 1985), the angle required for *O. serratus* to remove the librigenae and exit the exuvia would have applied more pressure to the anterior genal spine margins (Fig. 5). This approach to moulting likely resulted in the inflqated frequency of genal spine injuries in *O. serratus* (Fig. 1).

**Figure 5.**
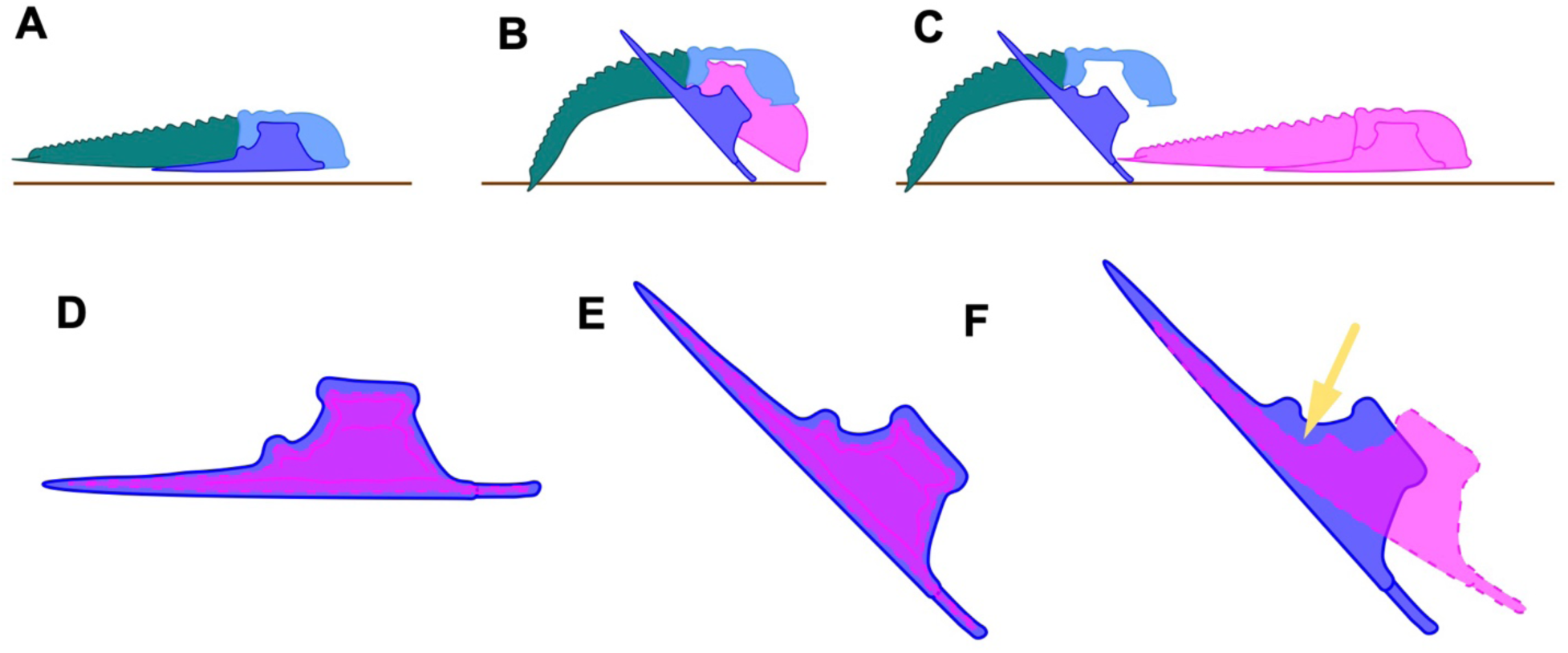
Moulting injuries in *Olenoides serratus*. (A) Pre-moulting individual. (B) Individual opening facial sutures by pressing into the substrate and beginning to exit the exuvia. (C) Individual exits exuvia. (D) Magnification of new moult underneath the librigenae prior to moutling. (E) Librigenae angles ventrally as facial sutures open in B. (F) The genal spine is injured by bending to exit the exuvia. A – C modified from Sam Gon III (https://www.trilobites.info/).

The likelihood of incurring genal spine injuries from moulting may depend on the size, robustness and function of the structure. It is possible that in trilobite species with more gracile genal spines, the new moult may be more flexible preventing injury. Whereas the genal spines are robust, they might better withstand the force applied from the angle (Fig. 5F). Two specimens of *M. splendens* are known with malformed cephalic spines (Whittington, 1971), possibly resulting from the moulting process as in *Olenoides serratus*. One specimen of *Marella splendens* from the Burgess Shale is preserved in the process of the new moult escaping the exuvia (García-Bellido and Collins, 2004). The cephalic spines are bent posteriorly acutely towards the axial margin of the exuvia. Unlike trilobites, *M. splendens* lacked a biomineralized exoskelton and the newly moulted individual was likely more flexible. Injuries to the librigenae including the palpebral lobe may have been caused by moulting (Šnajdr, 1978). Although librigenal injuries are generally rare (Owen, 1985) (Fig. 4), they often result from moulting events where the force to open the first suture or from exiting the exuvia (Fig. 5).

Although moulting mechanisms are well studied in trilobites based on specimen preservation and suture morphology (Daley and Drage, 2016), the dangers of this process which result in injuries and even death in modern euarthropods are not well understood. Injuries of spines in the trunk have been suggested to result from moulting, but a mechanism explaining this has not been suggested (Owen, 1985; Conway Morris and Jenkins, 1985; Bicknell and Pates, 2020). Our data on *Olenoides serratus* reveal an unusual pattern of injury in a trilobite and provide evidence for how these were incurred through moulting. The tradeoff of a rigid exoskeleton to provide protection against predators and the dangers of moulting said structure has been balanced throughout euarthropod evolution.

## Acknowledgements

We thank Jean-Bernard Caron and Maryam Akrami (Royal Ontario Museum, Toronto, Ontario, Canada), Michelle Coyne (Geological Survey of Canada, Ottawa, Ontario, Canada), Doug Erwin, Mark Florence and Gene Hunt (Smithsonian Institution, Washington D.C., USA), Omar Rafael Regalado Fernández and Olaf Vogel (Naturmuseum Senckenberg, Frankfurt, Germany) for facilitating access to specimens. This research was funded by a MAT Program Postdoctoral Fellowship and an Australian Research Council grant (DE250100256) to R.D.C.B.

**Table 1.**
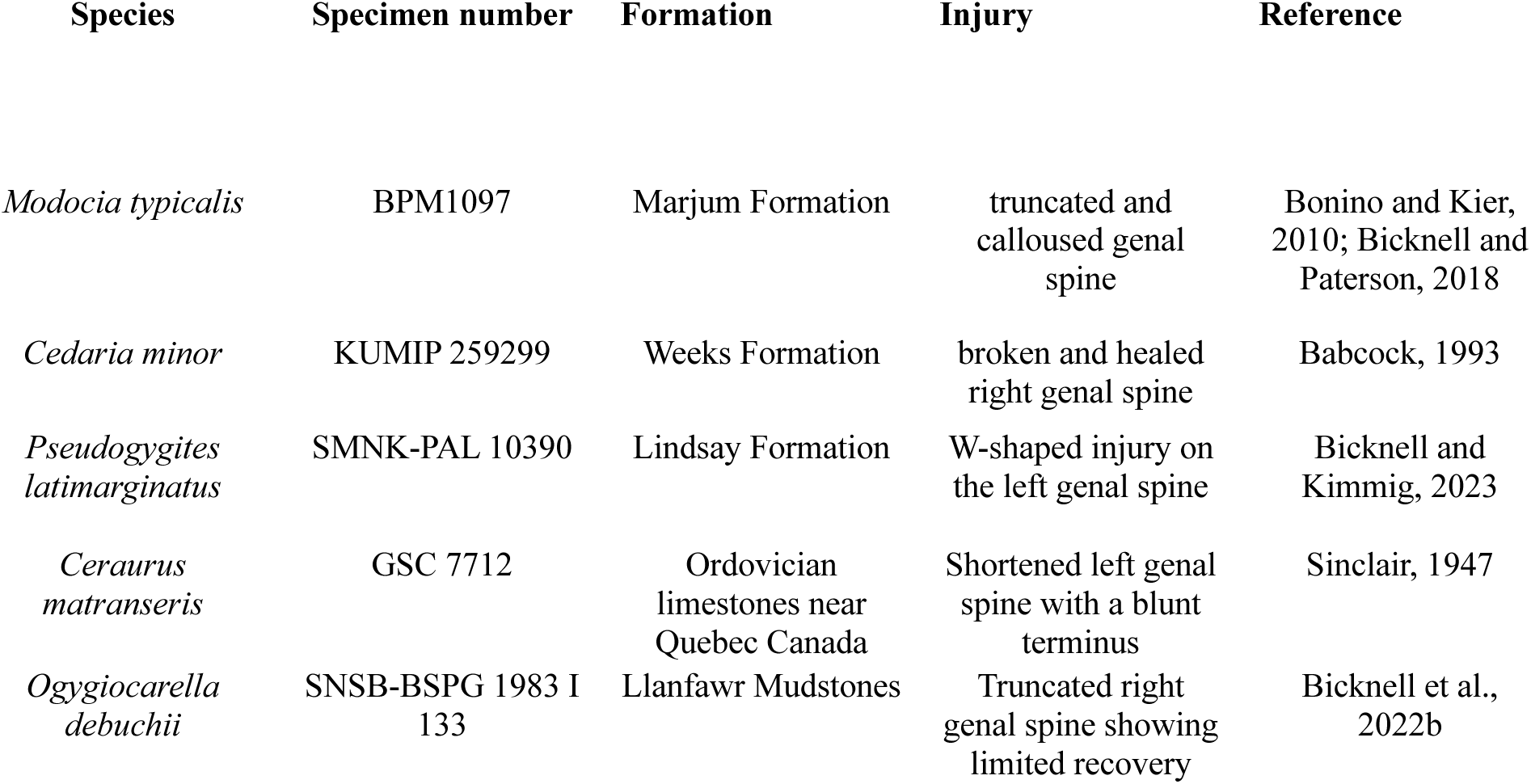
Known genal spine injuries in trilobites.

## Supplemental Information

**Supplemental Table 1. Institutional abbreviations.**

**Supplemental Table 2. Specimen measurements and occurrence of injuries.**

**Supplemental Table 3. Distribution of injuries in trilobites over the exoskeleton.**

**Supplemental Figure 1.**
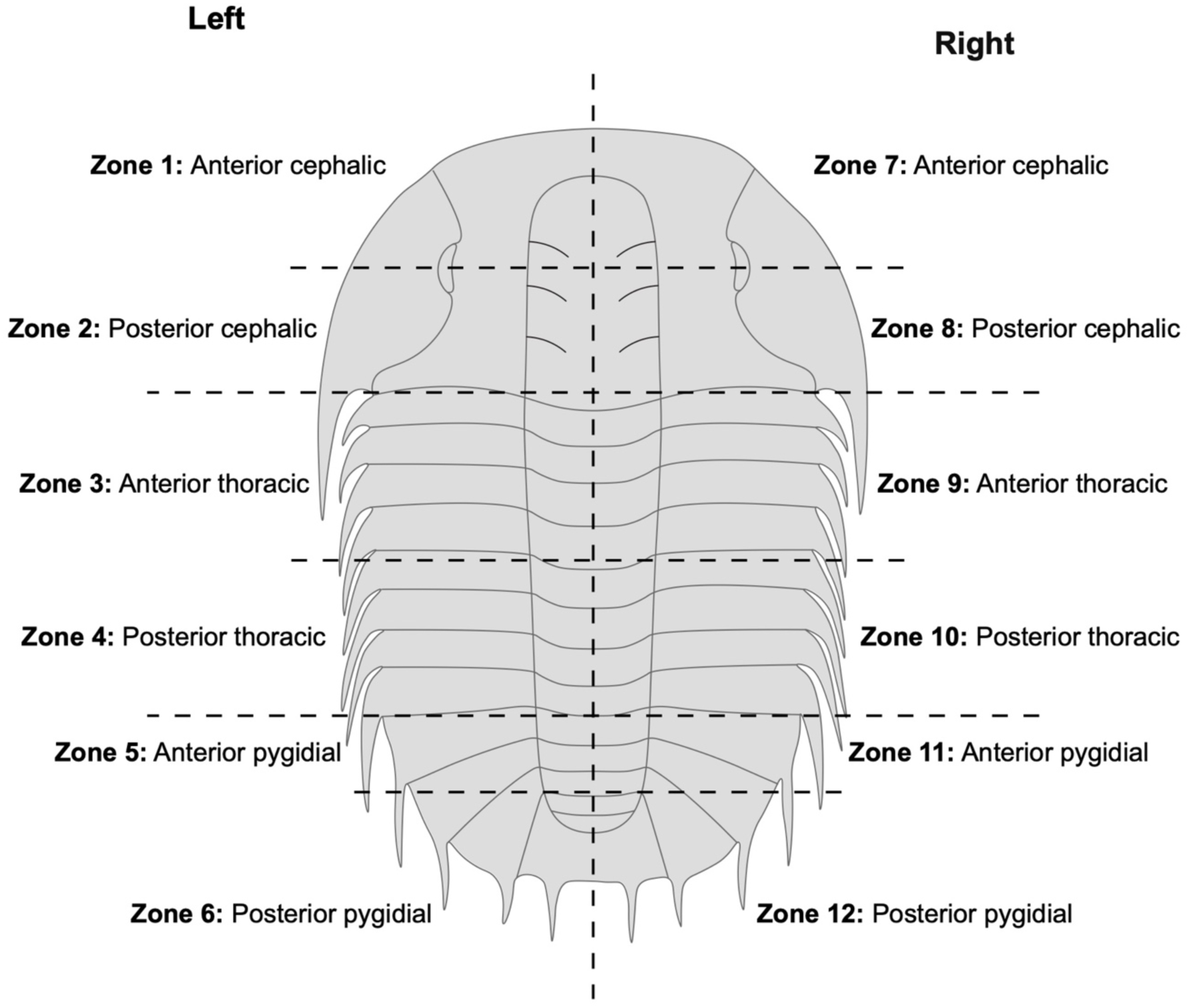
Zonation of the exoskeleton to evaluate distribution of injuries.

**Supplemental Figure 2.**
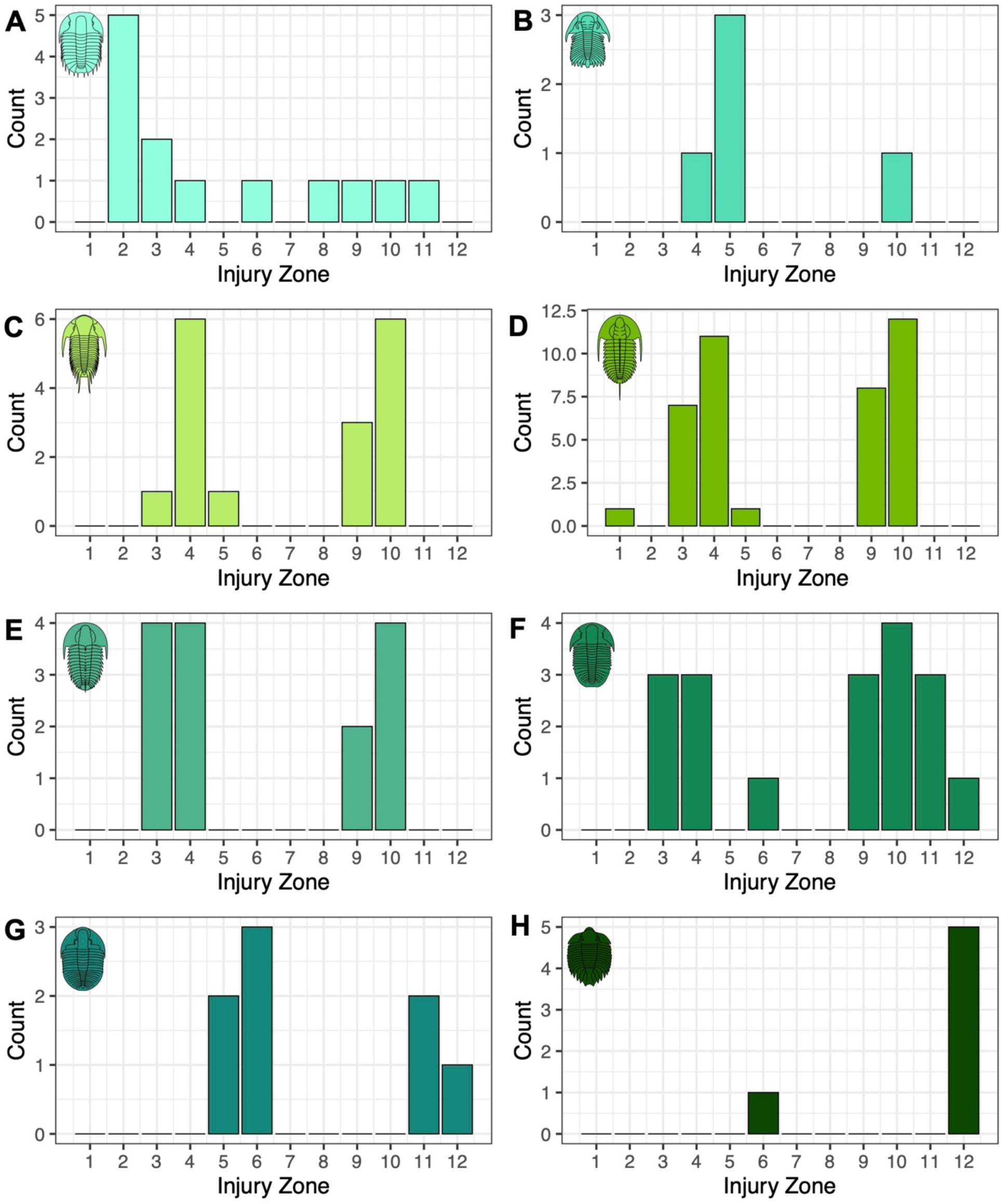
Histograms by species of number of injuries in each exoskeleton zone. (A) *Olenoides serratus*, this study. (B) *Eccaparadoxides pradoanus.* (C) *Paradoxides davidis trapezopyge*. (D) *Redlichia takooensis*. (E) *Redlichia rex*. (F) *Ogygopsis klotzi.* (G) *Ogygiocarella debuchii.* (H) *Arctinurus*. See Supplemental Figure 1 for exoskeleton zones and Supplemental Table 3 for data.

**Supplemental Figure 3.**
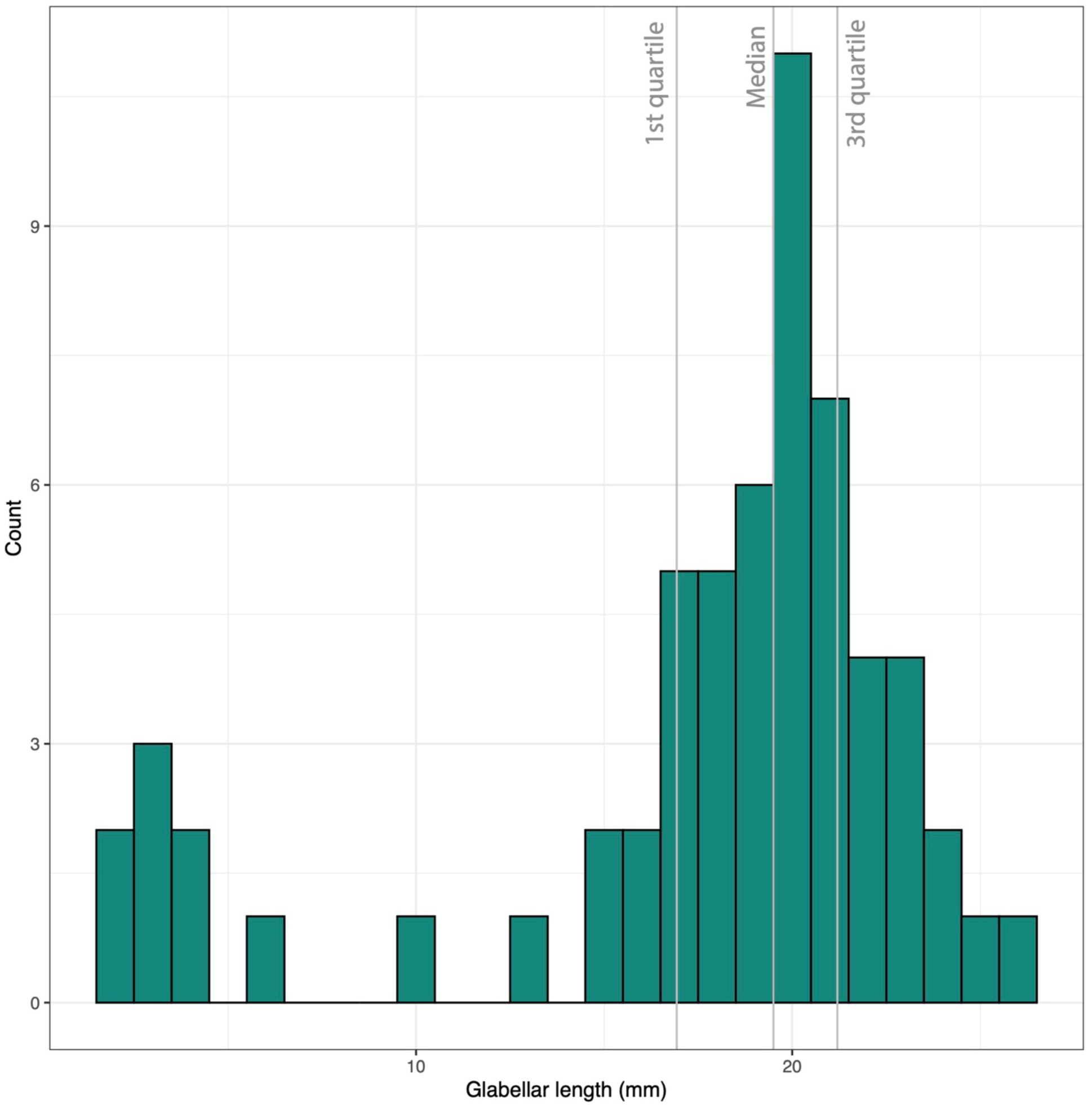
Histogram of glabellar length in *Olenoides serratus*.

